# Heat-stress triggers MAPK crosstalk to turn on the hyper-osmotic response pathway

**DOI:** 10.1101/374918

**Authors:** Paula Dunayevich, Rodrigo Baltanás, José Clemente, Alicia Couto, Daiana Sapochnik, Alejandro Colman-Lerner

## Abstract

Cells make decisions based on a combination of external and internal signals. In yeast, the high osmolarity response (HOG) is a mitogen-activated protein kinase (MAPK) pathway that responds to a variety of stimuli, and it is central to the general stress response. Here we studied the effect of heat-stress (HS) on HOG. Using live-cell reporters and genetics, we show that HS promotes Hog1 phosphorylation and gene expression, exclusively via the Sln1 phosphorelay branch, and that the strength of the activation is larger in yeast adapted to high external osmolarity. HS stimulation of HOG is indirect. First, we found that it depends on the operation of a second MAPK pathway, the cell-wall integrity (CWI), a well-known mediator of HS. Second, we show that HS causes glycerol loss via the channel Fps1, and that strictly requires the CWI MAPK Slt2. Third, blocking glycerol efflux also blocks HOG activation, strongly suggesting that it is the resulting loss of turgor by the loss of the accompanying water what causes HOG stimulation. Thus, taken together, our findings highlight a central role for Fps1, and the metabolism of glycerol, in the communication between the yeast MAPK pathways, essential for survival and reproduction in changing environments.

## Introduction

Sensing and responding to the environment is critical for survival. In eukaryotic cells, mitogen-activated protein kinase (MAPK) pathways mediate the response to many different stimuli. The budding yeast *S. cerevisiae* contains five such pathways ^1^. Two of them, the High Osmolarity Glycerol (HOG) ^2^ and the Cell Wall Integrity (CWI) ^3^ pathways are activated by a variety of environmental changes/conditions. The other three, the pheromone response ^4^, the filamentation ^5^ and the sporulation/meiosis pathways ^6^ control cell fate decisions in response to mating pheromone or nutritional conditions.

HOG responds to many stimuli, including acetic acid ^7^, arsenite ^8^, cold shock ^9^, and heat shock ^10^. The best characterized stimulus is a high osmolarity shock. An increase in external osmolarity causes loss of turgor pressure and cell volume, what triggers a homeostatic response leading to the accumulation of a compatible osmolyte, which in glucose containing medium is glycerol. Two signaling branches, known as Sln1 and Sho1 after their sensors, converge to ultimately activate the MAPKK Pbs2, which in turn activates the p38-like MAPK Hog1 ^11^ (Fig. 1A). In the Sln1 branch, this membrane sensor transduces the signal via Ypd1 and Ssk1 by a phosphorelay mechanism to the partly redundant MAPKKKs Ssk2 and Ssk22 ^12^, which phosphorylate Pbs2. In the other branch, the mucin-like sensors Msb2 and Hrk1 activate Sho1, which recruits a complex to the plasma membrane that includes Cdc42, promoting the activation of Ste20. Ste20 initiates the MAPK cascade by activating Ste11, which in turn phosphorylates Pbs2 ^13^. Once active, Hog1 stops glycerol efflux by phosphorylating regulators of the Fps1 channel, Rgc1 and Rgc2/Ask10. When phosphorylated, these proteins detach from the channel, which in turn closes ^14,15^. Hog1 also increases metabolic flow towards glycerol by up-regulating Pfk2 (a subunit of phosphofructokinase), inhibiting Ypk1, an inhibitor of Gpd1, the first step in the reduction of DHAP into glycerol ^16^ and blocking Tdh1,2 and 3, which divert the flow towards respiration. Phosphorylated Hog1 translocates to the nucleus where it associates with transcription factors like Hot1 ^17^ to induce a large number of genes ^18^, including some encoding enzymes and transporters required for glycerol accumulation ^19^.

**Figure 1.**
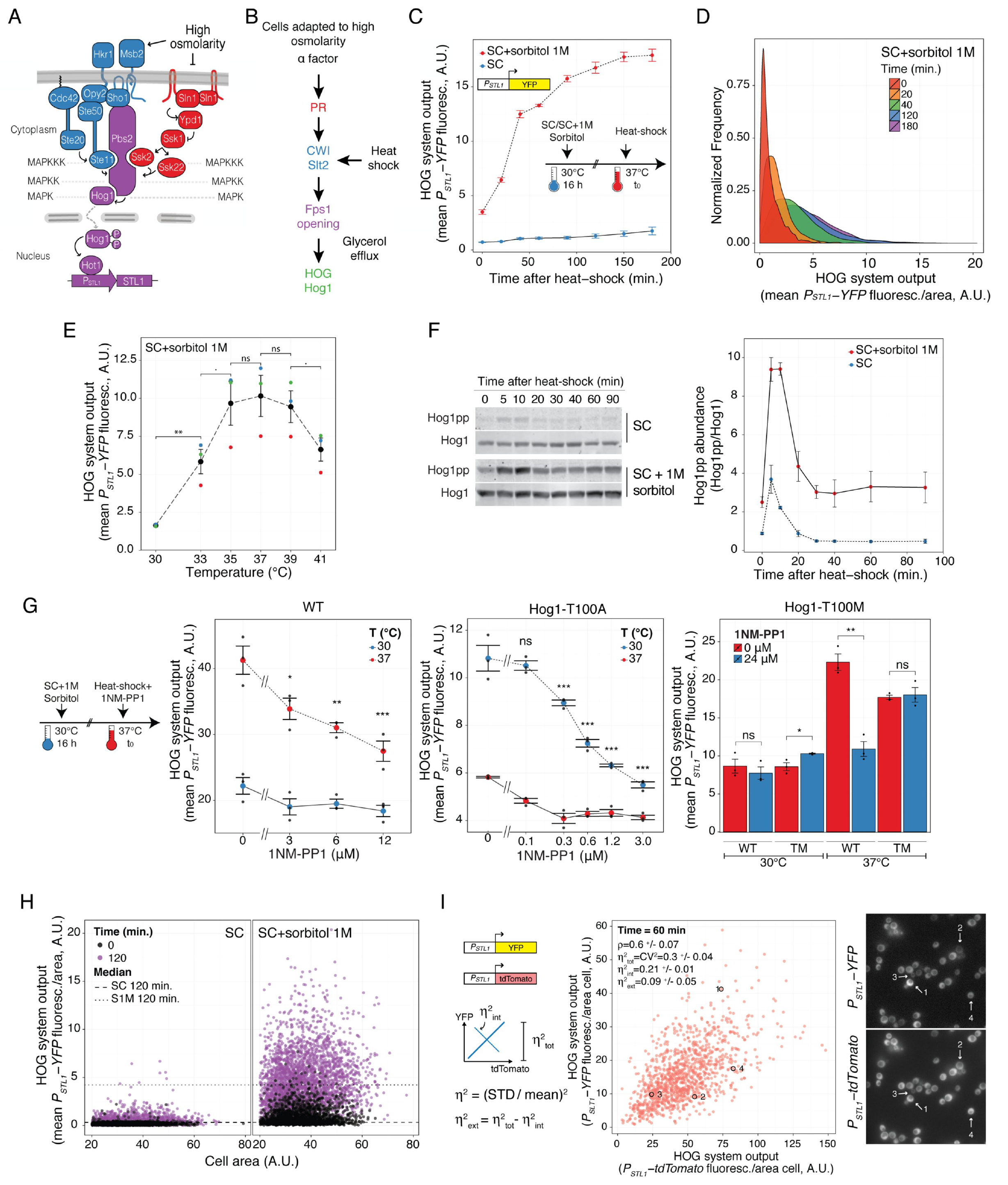
Heat-shock stimulates the HOG pathway. (A) The HOG pathway with its two branches (SHO1 in blue and SLN1 in red) converge on Pbs2, which phosphorylates and thus activates the MAPK Hog1. Active Hog1 translocates to the nucleus where it induces gene expression via the transcription factors Sko1, Skp1 (not shown), and Hot1. (B) Mating pheromone stimulates HOG indirectly. Pheromone activates the Cell Wall Integrity pathway, which in turn causes loss of glycerol through the aqua-glyceroporin Fps1, leading to reduced turgor, and causing HOG activation. We hypothesize that heat-shock (HS) stimulates HOG via this route as well. This activating pathway is amplified in cells adapted to high osmolarity medium. (C) HOG transcriptional reporter dynamics following a shift from 30°C to 37°C. Here, and in all similar charts elsewhere, plots of HOG system output (population average of the total YFP accumulated in each cell) vs. time correspond to the mean of three or more independent replicates ± SEM. (D) Histograms of the 1M sorbitol data as in C. (E) HOG activation by heat shock at the indicated temperatures shows a maximum at 37°C. Plot of HOG system output after 1 h of temperature shift corresponds to the mean of three or more independent replicates ± SEM. Colors correspond to replicate experiments. (F) HOG MAPK activation dynamics following a shift from 30°C to 37°C. Hog1 phosphorylation was assayed by immunoblotting. Left. Representative blot. Right. Here, and in all similar immunoblot quantification charts elsewhere, plot corresponds to the mean of three or more independent replicates ± SEM. Uncropped blot in Fig. S2. (G) Hog1 mediates the activation of the *P_STL1_-YFP* reporter by heat-shock. Left. WT Hog1 is naturally sensitive to the inhibitor 1NM-PP1: increasing concentrations of 1NM-PP1 progressively reduce heat shock stimulation of reporter protein. Middle. The Hog1-T100A mutant is hypomorph and supersensitive to 1NM-PP1. Right. The Hog1-T100M mutant is insensitive to 1NM-PP1, indicating that the effect observed in the WT is due to inhibition of Hog1 and not of another naturally sensitive kinase. Here, and in all subsequent bar charts elsewhere, unless otherwise indicated, HOG system output was measured after 2 h of temperature shift. Values correspond to the mean of three or more independent replicates ± SEM. Statistical comparisons of left and middle plots are against the 0 μM 1NM-PP1 at 37°C. (H) Scatter plot comparing data at 0 vs. 120 min after temperature shift of cells grown in SC or SC + 1M sorbitol. Plot shows HOG system output/area vs. area of individual cells. (I) Cell to cell variability in HOG activation. Strains with two identical *STL1* promoters driving YFP or tdTomato grown in 1M sorbitol medium after 1h shift from 30°C to 37°C. Plot shows YFP vs tdTomato protein reporters. ρ indicates Pearson correlation coefficient. Total variability is measured using η^2^_tot_ ((STD/mean)^2^); intrinsic noise η^2^_int_ (STD^2^_(YFP/<YFP> - tdTomato/<tdTomato>)_) and η^2^_ext_ (η^2^_tot_ - η^2^_int_) (see Methods). Images show a representative field. Numbers mark selected cells from the plot. White bar corresponds to 5 μm.

In contrast, a low osmolarity shock results in the aperture of Fps1 which allows glycerol, accompanied by water, to leave the cell ^20^. Therefore, the excessive pressure is alleviated. Opening of Fps1 seems to depend on the CWI pathway ^2^. The CWI is activated by many stimuli, including cell-wall binding compounds (like congo red and calcofluor white), heat shock and cell-wall remodeling events during the mating response ^3^. CWI signaling has a main pathway composed of several mucin-like proteins that operate as sensors connecting the plasma membrane and cell wall, and which display varying specificity for different stimuli. These sensors (Wsc1,2,3, Mid2, Mtl1) recruit the guanine nucleotide exchange factors Rom1 and Rom2, which activate the small G-protein Rho1. Rho1-GTP then activates the kinase Pkc1, which initiates the MAPK cascade by phosphorylating the MAPKKK Bck1. Bck1 stimulates the MAPKKs Mkk1 and Mkk2, which then activate the MAPK Slt2. Slt2, via transcriptional and post-transcriptional actions, causes remodeling and strengthening of the cell wall. Full activation of Slt2 is achieved, besides the activity of this main pathway, by parallel inputs downstream of Rho1, either to Pkc1, Bck1 or directly to Mkk1/2, depending on each particular stimulus ^21^. In particular, activation of Slt2 by heat-stress requires the main pathway and extra input to Mkk1,2, possibly from Cbk1 and Bck2 ^22^.

Thus, while both HOG and CWI are activated by acute changes in external osmolarity, they each respond to osmotic shocks in opposite directions. Nevertheless, other types of stimuli can activate both pathways ^23^. One example is heat stress, which elicits a complex response that involves a plethora of simultaneous actions by the cell ^24^. Among these responses, it activates both HOG ^10^ and CWI ^25^, although with different dynamics. While Hog1 shows a transient phosphorylation peak at around 5 minutes, Slt2 is slowly phosphorylated reaching a maximum at around 30 minutes.

In addition, we have previously shown that the HOG pathway is activated during the mating response in high-osmolarity ^26^. The mechanism that leads to this cross-talk involves the activation of the CWI MAPK Slt2 by the cell-shape changes associated to the mating projection formation, followed by the aperture of Fps1 (Fig. 1B). The resulting release of internal glycerol and loss of turgor pressure activates the Sln1 branch of HOG, and this leads to increased production of glycerol. This mechanism of cross-talk results in a high glycerol turn-over rate that allows fine and fast tuning of internal pressure within the yeast cell during mating ^26^, a pre-condition for fusion ^27^.

If the mating response is capable of activating HOG via the CWI pathway, it is likely that other CWI activating inputs would lead to a similar fate. This rationale might expand the role of the HOG pathway to include many inputs previously not thought to be associated with it. Here we show that both heat shock and the cell-wall disturbing agent congo red can cause HOG activation, which is strongly amplified when yeast are grown in high osmolarity medium. Similar to the cross-talk from the mating pheromone response, it involves the MAPK Slt2, the glycerol channel Fps1 and the Sln1 branch of HOG.

## Results

### Heat shock activates HOG in yeast adapted to high osmolarity

Our previous studies on the crosstalk between MAPK pathways indicated that activation of the CWI by mating pheromone causes HOG activation ^26^, especially noticeable in yeast pre-adapted to high osmolarity medium. Here we asked if CWI activation was sufficient for HOG activation (Fig. 1B). To test this idea, we stimulated the CWI by mild heat shock (shifting the culture from 30°C to 37°C) ^3^. We monitored HOG activation in individual cells using a transcriptional reporter (*P_STL1_-YFP*) (Fig. 1C). In SC medium, there was no detectable reporter induction. In contrast, in yeast grown in SC supplemented with 1M sorbitol, there was a fast increase in YFP synthesis. Accumulation of fluorescent protein approached steady-state after three hours, indicating that around this time, the higher reporter expression was compensated by dilution by cell division. Induction by heat-shock involved the whole population, as evidenced by an overall shift over time in the unimodal distribution of accumulated reporter (Fig. 1D). We detected a similar *P_STL1_-YFP* induction by heat shock after adapting cells to 0.5M NaCl (Fig. S1), indicating that it is the high external osmolarity and not sorbitol per se what is required for HOG stimulation.

We tested various temperature shifts and found maximum expression of the HOG reporter at 37°C. Higher temperatures resulted in a lower induction (Fig 1E). This could be because above 37°C, yeast might accumulate trehalose instead of glycerol ^28,29^, a process that is independent of HOG.

To confirm that the MAPK Hog1 was activated by heat-shock, we examined Hog1 phosphorylation by western blot (Fig. 1F and S2) after the temperature shift. In SC, as previously reported ^10^, there was a small, fast and transient phosphorylation of Hog1, with a peak at around 5 minutes, after which phosphorylated Hog1 dropped, reaching basal levels 30 minutes after heat-shock. In medium supplemented with 1M sorbitol, Hog1 phosphorylation dynamics in response to heat-shock was similar, but amplified: basal phospho-Hog1 was higher, with a higher peak signal.

To verify that reporter induction depended on Hog1 activity, we used the kinase inhibitor 1NM-PP1, an adenine analog. In most kinases, it is necessary to mutate the gatekeeper amino acid in the ATP binding pocket to a smaller residue to allow 1NM-PP1 to fit inside ^30^. However, we noted that Hog1 possesses threonine at this location, a relatively small amino acid, which might allow inhibition of the WT form (out of the 129 yeast kinases, 14 have Thr and 3 have Ala at this location, the rest have bigger residues). Indeed, 12μM inhibitor largely blocked reporter induction in response to a 0.8M sorbitol hyperosmotic shock (Fig. S3). This same concentration reduced reporter induction in response to heat-shock in yeast adapted to 1M sorbitol (Fig 1G, left), and 24 μM almost completely blocked it (Fig 1G, right). To control that the effect of the inhibitor was specific to Hog1, we constructed strains expressing Hog1-as2 (Hog1-T100A), which are more sensitive to 1NM-PP1 ^31^. This mutant is already hypomorph, exhibiting a reduced activation in response to heat shock in the absence of inhibitor (Fig. S3). Addition of 3μM of 1NM-PP1 blocked induction (Fig. 1G, middle). As a further control, we made a new mutant, Hog1-T100M, which has methionine in the gatekeeper position, a bigger amino acid. This mutant was resistant to the inhibitor both in response to hyperosmotic shock (Fig. S3) and to heat shock (Fig. 1G, right). Together, these results confirm that *P_STL1_-YFP* expression in response to heat shock requires Hog1 activity.

Taken together, these results indicated that heat-shock, stimulates the HOG pathway in cells already adapted to high external osmolarity.

### Single-cell analysis reveals large noise during heat-shock activation of HOG

The transient activation of Hog1 in SC did not result in noticeable reporter expression at the population level (Fig. 1C). However, further inspection of individual cells revealed that a fraction of less than 0.5% of SC grown yeast did induce the HOG reporter to levels comparable to those adapted to 1M sorbitol (note the cells with outlier abundance of YFP in Fig. 1H, left). This is reminiscent of the behavior of yeast exposed to small hyperosmolarity shocks (~ 0.1M NaCl), during which many cells fail to induce *SLT1* ^32^. Pelet et al. showed that expression of *STL1* requires a slow stochastic step of chromatin remodeling. When the stimulus is too short, the pathway turns off in a significant fraction of cells (measured by the exit of Hog1 from the nucleus) before the *STL1* locus had time to be remodeled and therefore became susceptible to induction. Due to the randomness in the timing of remodeling, this step introduced noise in the expression of *STL1* ^32^. Thus, we suspect that the reason for the lack of induction in SC is that, as during small hyperosmolarity shocks, Hog1 phosphorylation peak is either too short or too low in amplitude.

To assess the contribution of stochastic processes to the induction of *STL1* in response to temperature, we measured gene expression noise (also referred to as intrinsic noise) ^32-34^ 60 minutes after the temperature shift by comparing the induction of two identical *STL1* promoters driving the expression of YFP and tdTomato, respectively (Fig. 1I). The degree to which the signal from these promoters is decorrelated in the population is a measure of this noise. We found a mild correlation between YFP and tdTomato (ρ = 0.6 +/- 0.07), indicating a large contribution of gene expression noise. Indeed, this noise accounted for 70% of the overall cell to cell variability (η^2^_tot_ =0.3 +/- 0.04 vs. η^2^_int_ =0.21 +/- 0.01; η = STD/mean, see Methods). The remaining variability, known as extrinsic noise, corresponds the correlated variation between the YFP and tdTomato (η^2^_ext_ =0.09 +/- 0.05).

### The Sln1 branch mediates heat shock activation of HOG

Next, we tested which of the two HOG branches (Fig. 1A) mediates Hog1 activation by heat shock. Previous reports pointed to the Sho1 branch ^10^. However, deletion of *SHO1* did not prevent Hog1 phosphorylation after heat shock, neither in SC nor in yeast adapted to medium with 1M sorbitol (Fig. 2A and S4). In contrast, deletion of *SSK1* greatly diminished Hog1 phosphorylation, both in SC and in 1M sorbitol. Consistent with this result, the *P_STL1_-YFP* reporter was unaffected in *Δsho1* cells but was significantly reduced in *Δssk1*, *Δssk2 and Δssk22* cells (Fig. 2B). The double knockout *Δssk1Δsho1* showed an even lower reporter expression, suggesting a minor role for the Sho1 branch when the Sln1 one is inactivated.

**Figure 2.**
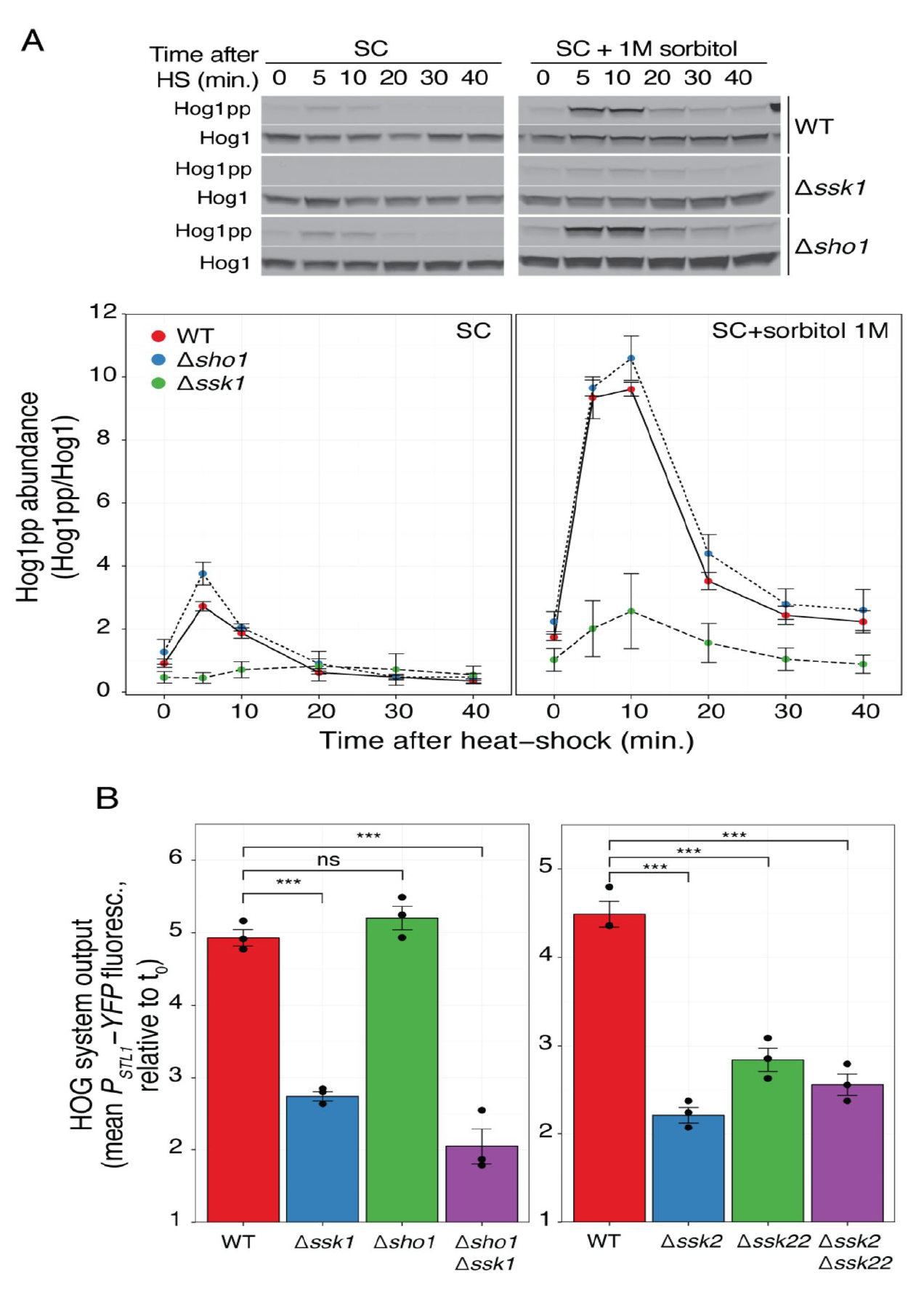
Heat-shock activation of HOG depends on the Sln1 branch. (A) HOG MAPK activation dynamics following a shift from 30°C to 37°C in branch mutants in SC or in SC + 1M sorbitol. Top. Representative blot. Uncropped blot in Fig. S4. (B) HOG transcriptional reporter in branch mutants measured 2 h after 1M sorbitol-adapted cells were shifted from 30°C to 37°C. Values correspond to the mean of three independent replicates ± SEM, relative to t_0_.

### Heat shock activation of HOG requires the CWI MAPK Slt2

Next, we tested the involvement of the CWI MAPK cascade. First, we confirmed that Slt2 is activated during heat-shock both in SC and in SC supplemented with 1M sorbitol by monitoring its phosphorylation (Fig. 3A and S5). Interestingly, Slt2 was already phosphorylated before the shock in both conditions, and it was further stimulated by heat-shock after an initial drop. Peak phosphorylation occurred between 15 and 20 minutes post stimulation, at a time in which Hog1 activation was already dropping to its lower steady-state. Next, we tested the effect of deleting *SLT2* on Hog1 phosphorylation (Fig. 3B and S2) and reporter induction (Fig. 3C). For both measures, the response of the *Δslt2* strain adapted to 1M sorbitol was greatly reduced. Single cell analysis showed that the remaining activity observed at the population average level was not due to a small fraction of active cells (Fig. 3D). It is noteworthy that the initial phosphorylation peak of Hog1 is diminished in *Δslt2* (both in yeast grown in SC or adapted to 1M sorbitol), since it indicates that the CWI plays a critical role in HOG activation from the start of the heat shock response, despite the fact that peak Slt2 phosphorylation occurs much later than Hog1’s peak.

**Figure 3.**
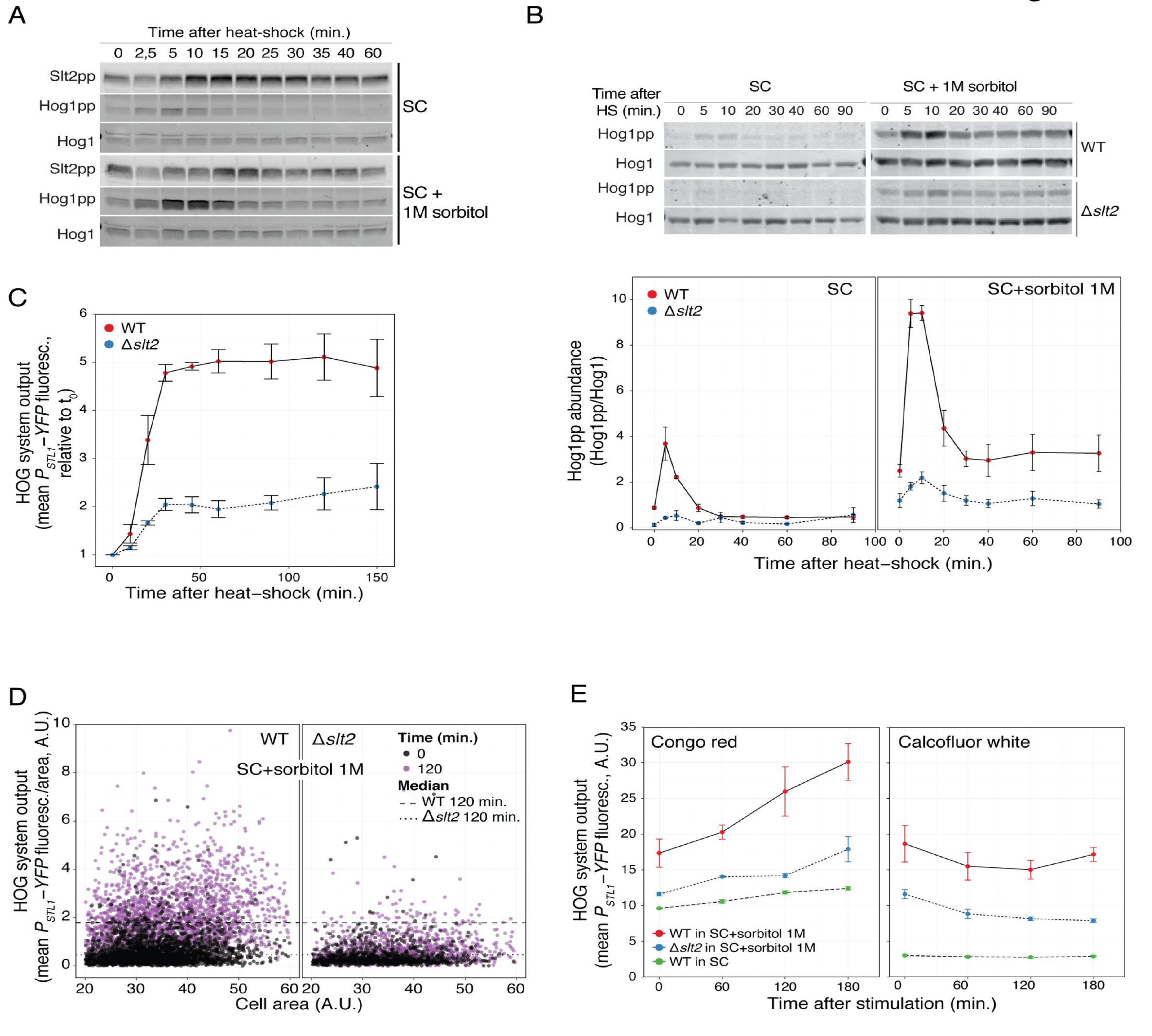
Heat-shock activation of HOG depends on the Cell-Wall Integrity pathway. (A) CWI MAPK activation dynamics following a shift from 30°C to 37°C in cells grown in SC or in SC+1M sorbitol. Slt2 phosphorylation was assayed by immunoblotting. Uncropped blot in Fig. S5. (B) HOG MAPK activation dynamics in WT and *Δslt2* cells grown in SC or in SC+1M sorbitol following a shift from 30°C to 37°C. Top: representative blot. Uncropped blot in Fig. S2. (C) HOG transcriptional reporter dynamics in WT and *Δslt2* following a shift from 30°C to 37°C. Values correspond to the mean of three independent replicates ± SEM, relative to t_0_. (D) Scatter plot comparing data at time 0 with 120 min after temperature shift. (E) HOG transcriptional reporter in stimulated with 50 μg/ml of congo red or 50 μg/ml of calcofluor white. Values correspond to the mean of three independent replicates ± SEM.

To this point, we have shown that two distinct inputs that cause Slt2 activation (shmoo formation during the mating response ^26^ and a temperature shock) result in HOG activation. We tested two other well-known CWI stimuli, the dyes congo red (50 μg/ml) and calcofluor white (50 μg/ml), which interfere with cell wall assembly by binding to chitin ^3^. At these concentrations, both drugs reduce yeast growth by 50% ^35^, and caused lysis of *Δslt2* cells in SC. Similar to heat-shock and pheromone, congo red induced *P_STL1_-YFP*, and it did so in manner dependent on high external osmolarity and the presence of the MAPK Slt2 (Fig. 3E). In contrast, calcofluor white failed to activate HOG, despite being able to activate Slt2. It may be that calcofluor white has extra effects that block either HOG or the connection between Slt2 and HOG (see Discussion).

### Loss of glycerol is essential for HOG activation

Our working model for the mechanism of HOG activation is centered around the glycerol-channel Fps1, with efflux of glycerol via this path as the connecting link between the CWI and HOG. To test the model, we deleted *FPS1* or its positive regulators *RGC1* and *RGC2*. Both in *Δfps1* and *Δrgc2*, as well as in the double knockout *Δrgc1Δrgc2*, induction of the HOG reporter was greatly reduced (Fig. 4A and S6). This is consistent with a crucial role of Fps1 channel in HOG activation. However, the single knockout *Δrgc1* showed only a partial reduction in reporter expression, indicating that this Fps1 regulator is not essential. This is interesting, since Rgc1 is required to activate HOG in response to mating pheromone ^26^ (Fig. S7), an indication that the pheromone and heat-shock mechanisms of HOG activation are not identical.

**Figure 4.**
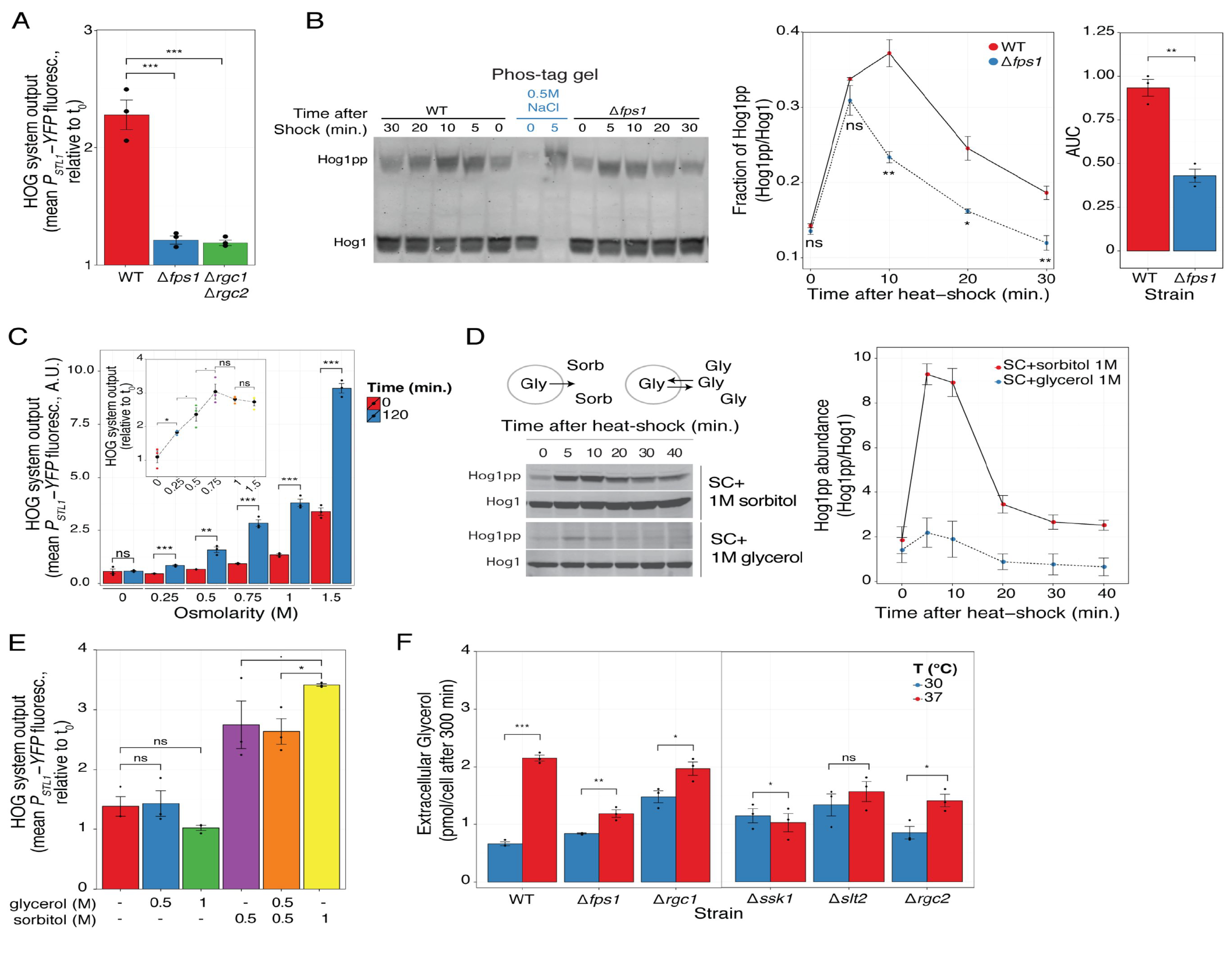
Heat-shock activation of HOG involves efflux of glycerol. (A) HOG transcriptional reporter in WT, *Δfps1 and Δrgc1Δrgc2* measured 2 h after a shift from 30°C to 37°C. Values correspond to the mean of three independent replicates ± SEM, relative to t_0_. (B) Phos-tag gel showing HOG MAPK activation dynamics in WT and *Δfps1* grown in SC + 1M sorbitol following a shift from 30°C to 37°C. Left: representative blot that includes a hyper-osmolarity shock control (blue text). Middle. Fraction of phosphorylated Hog1. Right. Mean area under the curve (A.U.C.) of phosphorylated Hog1 ± SEM. Statistical comparisons are WT vs *Δfps1*. Uncropped blot in Fig. S9. (C) HOG transcriptional reporter in WT adapted to the indicated osmolarities at 30°C and then transferred to 37°C. Inset shows the increase in reporter due to the temperature shift relative to t_0_. (D) HOG MAPK activation dynamics in WT adapted to 1M sorbitol or 1M glycerol, and then transferred to 37°C. Diagram. In medium with sorbitol, when Fps1 opens, glycerol will be lost from the cell. However, in medium with glycerol, even if Fps1 opens, the net glycerol flux should be zero. Left. Representative blot. (E) HOG transcriptional reporter in WT adapted to the indicated mixtures of glycerol and sorbitol at 30°C and then transferred to 37°C. Values correspond to the mean of three independent replicates ± SEM, relative to t_0_. (F) Accumulated extracellular glycerol during 300 minutes of growth at 30°C or after transfer to 37°C. Plot shows the mean of pmols of glycerol/cell ± SEM.

Surprisingly, Hog1 phosphorylation was not eliminated in Δ*fps1* relative to WT (Fig. S8): the initial peak was similar, although phospho-Hog1 abundance dropped faster. To confirm this unexpected result, we used phos-tag polyacrylamide gels, which provide a way to separate the unphosphorylated from the phosphorylated forms of Hog1 efficiently ^36^, so that a single antibody against Hog1 may be used to detect all its forms, and therefore improving the quantification of the fraction of phosphorylated Hog1. Two results from these gels (Fig 4B and S9) are worth mentioning. First, heat-shock caused a mobility shift identical to a control hyperosmolarity shock, confirming that Hog1 is dually phosphorylated ^36^. Second, the dynamics of Hog1 phosphorylation in WT and *Δfps1* was very similar, confirming that Hog1 indeed becomes phosphorylated after heat shock in *Δfps1* yeast. As with the conventional polyacrylamide gels (Fig. S8), phosphorylation dropped to values similar to pre-shock and in *Δfps1*, it dropped faster. As a consequence, total Hog1 activity over the 30 minute experiment in *Δfps1* was 50% of that in WT cells (Fig. 4B right). These results indicate that the initial phosphorylation peak is largely independent of Fps1. They also suggest that for induction of gene expression, the initial phosphorylation peak is not enough. Instead, sustained phosphorylation, likely involving continued loss of glycerol via Fps1, might be necessary (see Discussion).

We had previously found that the degree of HOG stimulation by pheromone positively correlated with the external osmolarity of the medium used to grow the cells ^26^. Similarly, heat shock induced the HOG reporter more strongly at higher osmolarities (Fig. 4C) while a control reporter was not affected (Fig. S10). Although accumulation of reporter did not reach an absolute maximum at the highest osmolarity we tested (1.5M sorbitol), fold-change induction plateaued at 0.75M sorbitol (Fig. 4C inset). This dependence on external osmolarity suggested that loss of glycerol was at the core of HOG activation during heat shock. This is because at higher external osmolarities, cells need to accumulate glycerol to correspondingly higher concentrations. If heat shock causes glycerol loss, more will be lost at higher osmolarities (due to a larger chemical gradient), turgor will be reduced more, and HOG will be activated to a greater extent. To test the role of glycerol loss, we replaced the added sorbitol with glycerol, eliminating in this way the concentration gradient driving glycerol out. Thus, even if cells open Fps1, the net flux should be zero. This change resulted in Hog1 phosphorylation to the same levels observed in SC (Fig 4D and S11) and abolished reporter induction (Fig 4E and S12). We also tested a 0.5M sorbitol-0.5M glycerol mix, which has the same external osmolarity as 1M sorbitol but a chemical gradient equivalent to just 0.5M sorbitol. In this mix, HOG activation reached levels identical to just 0.5M sorbitol. These results support the role of glycerol loss as the driver for HOG activation by heat-shock (Fig 4E).

As a final test of our model, we determined if heat shock causes the loss of glycerol by measuring its extracellular accumulation after the shift to 37°C. This measure reflects the combined ability of cells to release and to synthesize glycerol. Indeed, WT cells subjected to heat shock accumulated glycerol to a larger extent than without the shock (Fig. 4G). In contrast, the effect of temperature shift was absent (*Δslt2* and *Δssk1*) or greatly reduced (*Δfps1 and Δrgc2*) in the strains that fail to activate HOG. Of note, the strain that had the largest residual ability to release glycerol while at 37°C, *Δrgc1*, is the strain that showed only a partial defect in HOG activation by heat stress. These results further confirm that glycerol loss plays a central role in HOG activation. Note that even though *Δfps1* cells lack the only glycerol channel, heat-shock did increase glycerol accumulation in these cells. This suggests that there is an alternative route for glycerol efflux, perhaps via passive diffusion through the plasma membrane, and it might explain why Hog1 is phosphorylated in this strain (Fig. 4B). According to our model, *Δssk1* cells should not show any defect in Fps1 aperture upon heat shock. Thus, the failure of this strain to accumulate glycerol perhaps reflects its inability to activate of HOG and in this way boost glycerol production in these conditions.

## Discussion

Here we have shown that stimulation of the CWI pathway causes HOG activation. Our data is consistent with the mechanism of activation involving the release of glycerol in a manner dependent on the MAPK Slt2 (Figure 5). The exit of glycerol is accompanied by water and thus causes loss of turgor pressure, leading to the activation of the HOG pathway. Activated Hog1 stimulates glycerol production so that cells can reestablish their turgor pressure. Continued glycerol efflux maintains HOG active. The entire process is amplified when yeast grow in high external osmolarity, since the gradient driving glycerol out is greater under such conditions.

**Figure 5.**
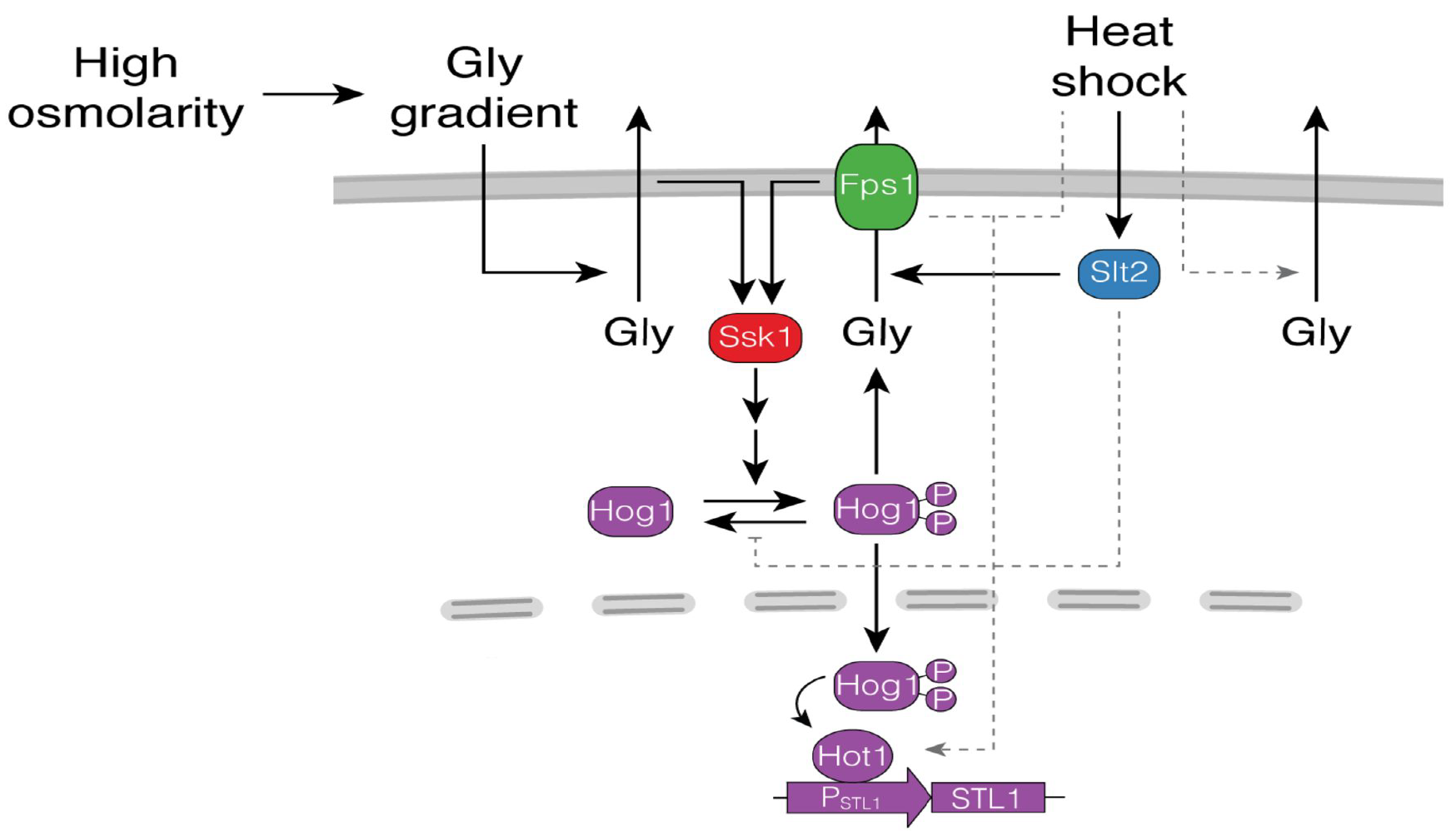
Model of HOG activation by heat-shock. Heat-shock stimulates the Slt2 MAPK, which in turn stimulates glycerol efflux possibly by causing the opening of the Fps1 channel. Significant efflux requires a large chemical gradient, which happens when yeast grow in high osmolarity medium. Glycerol efflux causes reduced turgor pressure what in turn stimulates the Sln1 branch (Ssk1) of HOG. Activated Hog1 induces gene expression (*P_STL1_-YFP*) and glycerol synthesis, which maintains a high chemical gradient, closing the circle. Our evidence also supports separate mechanisms (in dotted lines) by which a) Slt2 increases Hog1 activity, perhaps by inhibiting Hog1 phosphatases, b) Fps1 regulates Hog1-dependent gene expression downstream of Hog1 phosphorylation, and c) heat-shock causes glycerol efflux in a Fps1-independent manner allowing marginal activation of HOG even in *Δfps1* and *Δslt2*.

Upon heat-stress, Hog1 is phosphorylated following a “peak and decline to a lower plateau” dynamics. In our W303, phosphorylation was dependent on the Sln1 branch, both in SC and in high osmolarity medium. This is in contrast to a previous report that showed exclusive dependence on the Sho1 branch, at least in SC ^10^. We have no explanation for this discrepancy other than possible differences between DF5 and W303 genetic backgrounds. Our data indicates that strong Hog1 phosphorylation depends on high external osmolarity, on glycerol gradient and on the activity of Slt2. In contrast, the shape of its dynamics is independent of both, since a small peak and decline is still detectable in *Δslt2* yeast grown in SC. The effect of external osmolarity may be safely attributed to the amplitude of the glycerol gradient, and not to external osmolarity per se, since using glycerol instead of sorbitol reduces Hog1 phosphorylation to the values observed in SC. Initially, we hypothesized that the role of Slt2 on Hog1 phosphorylation could be completely explained by assuming that Slt2 causes Fps1 aperture, triggering the above explained activation of the Sln1 branch. This idea was further supported by our observation that Slt2 was required for the heat-shock-dependent increase in accumulated extracellular glycerol. However, the observation that in *Δfps1* yeast, peak phosphorylation of Hog1 appeared almost as high as in WT yeast, indicated that Slt2 regulates Hog1 phosphorylation during heat-shock also by another mechanism, perhaps by inhibiting HOG pathway phosphatases, such as Ptp2, Ptp3 or Ptc1,2,3, all of which can target Hog1 ^37^. Such inhibition might be important, given that the HOG pathway has a significant basal activity, even in SC ^38^, which is further increased in yeast adapted to high osmolarity (evident at time zero in our western blots and reporter assays (Fig. 1)).

The behavior of *Δfps1* strain was surprising. This strain cannot stimulate *P_STL1_-YFP* expression in response to heat-shock, even though it exhibits robust Hog1 phosphorylation. This decoupling of phosphorylation and transcription has been described previously in the response to arsenite or antimony stress ^8^, although no mechanism was suggested. On the one hand, it is possible that the somewhat faster drop in phosphorylated Hog1 in *Δfps1* cells could be enough to prevent Hog1 dependent transcription. This is what happens in many cells exposed to a small hyperosmotic shock, where Hog1 returns to the cytoplasm before the *STL1* locus is induced ^32^. Alternatively, Fps1 might serve a second function, unrelated to glycerol release, required for Hog1 to stimulate transcription in this context (Figure 5).

We have stimulated the CWI MAPK Slt2 with four different external stimuli (mating pheromone ^26^, heat-stress, congo red and calcofluor white). All but calcofluor white stimulated the HOG pathway and all in a manner that depended on Slt2. Given that calcofluor white activates Slt2, the inability of this drug to activate HOG suggested it might have an extra effect that prevented *P_STL1_-YFP* induction. Global gene expression analysis provides some support for this idea, since the two drugs induce only partly overlapping gene expression profiles ^35^. In agreement with our result, Talemi et al. ^39^ also found that calcofluor white does not cause Hog1 phosphorylation. The other possibility, namely that congo red has an extra effect necessary to activate HOG, is less likely, since the two other CWI stimuli we used (mating pheromone and temperature) did activate HOG.

In conclusion, we found that heat-shock robustly stimulates HOG in a largely indirect fashion via the CWI. We suggest that Fps1 acts as a key regulatory node whereby different MAPK cascades converge to jointly adjust stress responses. The net stimulation of HOG by heat-shock varies depending on the actual environmental conditions, from low to high in accordance with the level of external osmolarity. The wealth of accumulated knowledge about MAPK cascades in yeast offers the opportunity to study how these networks of signaling evolve and to generate hypothesis and working models as to how similar networks operate in higher eukaryotes.

## Materials and methods

### Genetic and molecular biological methods

All nucleic acid manipulations and yeast methods were done as described ^40^.

### Yeast strains

Strains used in the paper appear in Table 1.

P_STL1_-YFP reporter strains derive from LD3342 ^26^, in turn derived from ACL379 (*can1∷P_HO_-CAN1 ho∷P_HO_-ADE2 bar1Δ ura3 ade2 leu2 trp1 his3*) ^34^, a W303-1a descendant. The strain has three reporters: a HOG-inducible reporter (*P_stl1_-YFP*) inserted in the *STL1* promoter; a mating pheromone-inducible reporter (*P_PRM1_-mCherry*) and a constitutive reporter, *P_BMH2_*-*CFP*, in *BHM2* locus.

We made *P_STL1_-tdTomato* reporter strains by inserting plasmid pJT3557 ^41^, a Yiplac128 bearing *P_STL1_-tdTomato*, in *STL1* locus by cutting it with NruI, into strain ACL3341, and selecting in - leucine medium.

We made deletions by PCR, as described ^42^.

To make the *HOG1-T100A* and *HOG1-T100M* containing strains, we made a PCR fragment covering the entire *HOG1* ORF with the T100A and T100M mutation in two steps. First, we generated two PCR fragments, using yeast genomic DNA as template, that overlap at the location of the desired mutation using the following oligos:

oHog1-prom-fw: ACCTCAAAGCGCTTCGTCATGG
oHog1-term-rev: TATTTATGAAAATTCCTCTTCGG
oHog1-TA-direct: TGGAAGATATATATTTTGTC**GCT**GAATTACAAGGAACAGATTTACATAGAC
oHog1-TA-reverse: GTAAATCTGTTCCTTGTAATTC**AGC**GACAAAATATATATCTTCCAATGGAGAAAG
oHog1-TM-direct: TGGAAGATATATATTTTGTC**ATG**GAATTACAAGGAACAGATTTACATAGAC
oHog1-TM-reverse: GTAAATCTGTTCCTTGTAATTC**CAT**GACAAAATATATATCTTCCAATGGAGAAAG

For Hog1-T100A (Hog1-T100M), we generated fragments using *oHog1-prom-fw* and *oHog1-TA-reverse (oHog1-TM-reverse)*, and *oHog1-term-rev* with *oHog1-TA-direct (oHog1-TM-direct)*. We then performed a second PCR with the external oligos *oHog1-prom-fw* and *oHog1-term-rev* using the two PCR products generated in the first step for each mutation as template. Finally, we transformed this PCR product into a *Δhog1∷kanMX* strain, YRB3801 ^26^, and selected transformants in YPD plates supplemented 0.5M NaCl (in which the parental *Δhog1* does not grow). We confirmed the resulting strains by sequencing.

### Single-cell microscopy methods

We cultured yeast in SC medium supplemented or not with NaCl, sorbitol and/or glycerol and allowed them to grow to exponential phase for at least 15 hours at 30°C with agitation. We used cultures with low absorbance (A_600nm_ between 0.05 and 0.3) to minimize cellular autofluorescence, especially in the vacuole ^34,43^. We took samples for microscopy and added cycloheximide (50 mg/ml) to stop further translation and allow full maturation of the reporter fluorescent proteins for at least 3 hs before imaging ^34^.

We imaged using an oil-immersion 60x PlanApo objective (NA = 1.4) in an Olympus IX81 inverted microscope equipped with CoolLED illumination, a motorized XYZ stage, and a CoolSnapHQ2 cooled charge-coupled device camera (Photometrix) in a 30°C room. We used MetaMorph 7.5 software (Universal Imaging Corporation) to image and control the microscope. We acquired a bright-field, mCherry or tdTomato, YFP, and/or CFP fluorescence images using filter sets 41004, 31044v2, and 41028 from Chroma Technologies Corp.

To extract quantitative information, we processed images with VCell-ID as previously described ^43,44^, and analyzed it with R (http://www.r-project.org) and the package Rcell (http://cran.r-project.org/web/packages/Rcell/index.html).

To calculate the transcriptional output (defined as the total corrected fluorescence signal in a cell from the corresponding reporter gene), we subtracted the background signal measured outside cells from the integrated total fluorescence for a given cell (the sum of the fluorescence values of all the pixels that VCell-ID associated with that cell). When indicated, we normalized this value to the area of the cells.

### Analysis of cell-to-cell variability

We performed the analysis as in Colman-Lerner et al. ^34^. We measured total variability as η^2^_tot_ (STD_(xFP)/<xFP>_), η^2^_int_, gene expression noise ("intrinsic noise" ^33^) as the variance in the difference of the normalized abundance of the two fluorescent proteins driven by identical copies of *P_STL1_* (STD^2^ _(YFP/<YFP> - tdTomato/<tdTomato>)_), and η^2^_ext_, extrinsic noise, as the difference between η^2^_tot_ and η^2^_int_.

We estimated the correlation ρ using the cor() function of R (Pearson method).

### Protein methods

We prepared samples essentially as described ^45^. Briefly, for each time point, we mixed 2ml of exponentially growing cells (A600 nm ≤ 0.6) with 165μL of 10M NaOH and 20μL of 100% 2-mercaptoethanol (premixed immediately before the experiment). We then incubated samples at room temperature for at least 5 min, then we added 300ml of 100% trichloro acetic acid (Sigma-Aldrich), followed by 10min on ice, and centrifugation at 16000 g for 2min at 4°C. We washed the pellet with 900μl of an ice-cold acetone solution (70% (v/v) acetone in 100mM tris-HCl, pH 6.8) by vortex, followed by re-centrifugation. We air-dried the pellet, and resuspended it in 50μl of resuspension buffer (100mM tris (pH 6.8), 3%) for 30min, switching between heating at 65°C and a Genie disruptor (Scientific Industries) (total time of disruptor 10min). Finally, we centrifuged samples for 10min at 16000g (at 4°C) to clear cellular debris and stored 40μl supernatants −80°C, until used in conventional PAGE or phos-tag gels.

For gel loading, we mixed 40μl of protein solutions with 10μl of SDS–polyacrylamide gel electrophoresis loading buffer 5X (2-mercaptoethanol 25%, 50% glycerol, 10% SDS, 150mM tris (pH 6.8), 0.002% bromophenol blue) pre-heated at 65°C, boiled for 5min, place on ice, and loaded on the gel.

We used 12% tris-HCl/glycine polyacrylamide gels (acrylamide 15%, Tris 375mM pH 8.8, 0,1% SDS, 0,1% APS, 0,004% TEMED), then transferred onto a 0.2-mm Immobilon-FL polyvinylidene difluoride membrane (IPFL00010, Millipore). We blocked membranes for at least 1h at room temperature in blocking buffer solution (TBS - 10mM Tris (pH 7.4), 150mM NaCl-, milk 5%, tween-20 0.05%). We detected phosphorylated MAPK Mpk1/Slt2 with a rabbit anti–phospho-p44/p42 at 1:1000 dilution (Cell Signaling Technology, #9101 L), pp-Hog1 with a mouse anti–phospho-p38 at 1:1000 dilution (Cell Signaling Technology, #9216 L) and total Hog1 with an anti-Hog1 (#SC-6815) at 1:1000 dilution (Santa Cruz Biotechnology Inc.) in blocking buffer solution.

We washed in TBS with 0.05% Tween and probed for 1h at room temperature with an 800 nm donkey anti-rabbit (#926-32213 IRDye 800CW), an 800nm donkey anti-mouse (#926-32212 IRDye 800CW), a 680nm donkey anti-goat (#926-68074 IRDye 680RD) fluorophore-conjugated secondary antibodies (Invitrogen) and a 680 nM donkey anti-rabbit (#926-68073 IRDye 680RD); all diluted 1:10,000 in blocking buffer solution.

Finally, we washed blots 3 times in TBS with Tween and once in TBS and then scanned them on a Li-Cor infrared imaging system (Li-Cor Biosciences). We quantified band intensities with the Odyssey Image Studio software (Li-Cor Biosciences) using the median intensity of pixels in a border around a shape for background correction.

For phos-tag gel electrophoresis, we used 8% polyacrylamide with 20μM phos-tag (Wako Chemical Industries #304-93521) following the protocol described previously ^36^, except for the transfer buffer; we used Tris-Glycine-Methanol, as described for conventional PAGE.

### Glycerol measurements

Extracellular glycerol was measured essentially as described ^26^. Yeast cells were grown overnight in SC medium supplemented with 1M sorbitol (SCS) at 30°C. For this we started with a diluted culture to obtain an A_600nm_ of 0.2 the following day. Next day, the culture was filtered (cellulose ester filtering discs of 0.45μm pore size and 25mm diameter; Millipore, catalog no. HAWP02500) in a “1225 manifold” (Millipore, catalog no. XX2702550) and resuspended in fresh SCS medium. This step is necessary to remove glycerol accumulated before the experiment. Next, we split this culture into two flasks, one for the 37°C heat shock and the other for 30°C control. After 300 minutes we filtered 5mL of cells (HAWP02500, Millipore filtered) and collected the flow through in 15mL falcon tubes. Finally, the concentration of glycerol was measured by high-pH anion exchange chromatography with pulse amperometric detection in an ICS-3000 chromatographic system (Dionex), as previously described ^46^. We used a CarboPac MA1 column (4 × 250mm, Dionex) and a CarboPac MA1 guard column (4 × 50mm, Dionex). An isocratic program with 200mM NaOH was used at a flow rate of 0.5ml/min and a loop of 20ml. The standard curve was measured between 0 and 4000ng of glycerol. Samples were diluted as necessary to obtain data in this range.

The following mathematical transformation was used to calculate the amount of glycerol produced *per cell*:

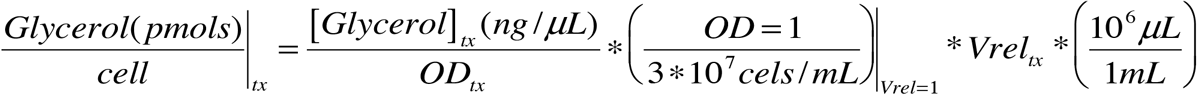

where Vrel_tx_ is the relative volume of cells at a certain time (tx) relative to the volume of cells at time 0.

In the same experiments, we collected samples and added cycloheximide to a final concentration of 100mg/ml; measured absorbance; and imaged cells to quantify cell volume and reporter expression.

### Statistical methods

For statistical significance determination of fluorescence transcriptional reporters (*P_STL1_-YFP*, *P_BMH2_-CFP*, *P_PRM1_-mCherry*), and western blot data, we used ANOVA in at least three biological replicates experiments. Error bar corresponds, unless otherwise indicated, to the standard error of the mean (SEM). Statistical significance is indicated as follows: “ns” p> 0.05, “·” p 0.1, *: p<0.05, **: 0.01, *** <0.001. In all data with transcriptional reporters, number of yeast was greater than 200.

**Table 1.**
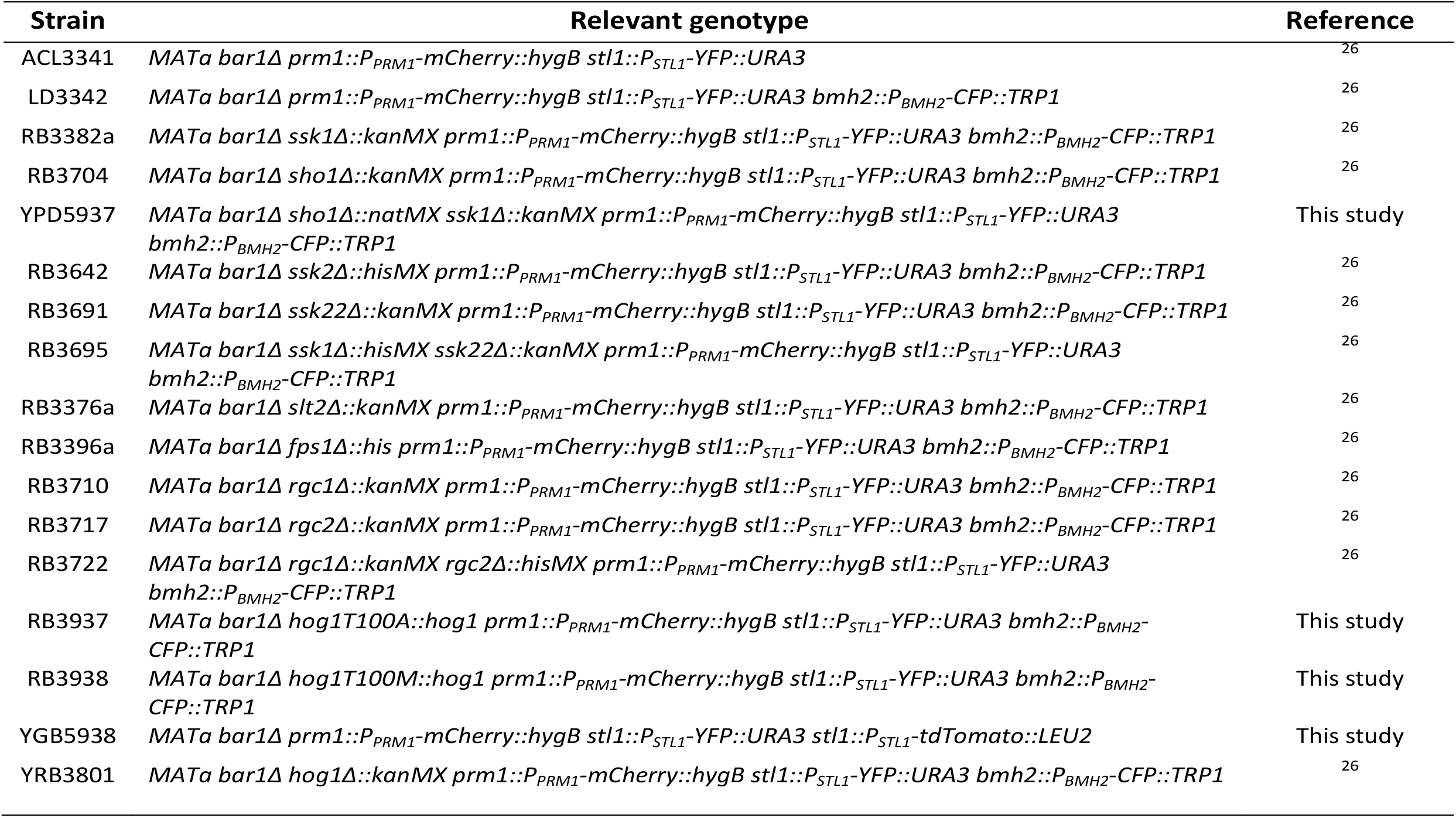
Yeast strains.

| Strain | Relevant genotype | Reference |
| --- | --- | --- |
| ACL3341 | *MATa bar1Δ prm1::P<sub>PRM1</sub>-mCherry::hygB stl1::P<sub>STL1</sub>-YFP::URA3* | 26 |
| LD3342 | *MATa bar1Δ prm1::P<sub>PRM1</sub>-mCherry::hygB stl1::P<sub>STL1</sub>-YFP::URA3 bmh2::P<sub>BMH2</sub>-CFP::TRP1* | 26 |
| RB3382a | *MATa bar1Δ ssk1Δ::kanMX prm1::P<sub>PRM1</sub>-mCherry::hygB stl1::P<sub>STL1</sub>-YFP::URA3 bmh2::P<sub>BMH2</sub>-CFP::TRP1* | 26 |
| RB3704 | *MATa bar1Δ sho1Δ::kanMX prm1::P<sub>PRM1</sub>-mCherry::hygB stl1::P<sub>STL1</sub>-YFP::URA3 bmh2::P<sub>BMH2</sub>-CFP::TRP1* | 26 |
| YPD5937 | *MATa bar1Δ sho1Δ::natMX ssk1Δ::kanMX prm1::P<sub>PRM1</sub>-mCherry::hygB stl1::P<sub>STL1</sub>-YFP::URA3 bmh2::P<sub>BMH2</sub>-CFP::TRP1* | This study |
| RB3642 | *MATa bar1Δ ssk2Δ::hisMX prm1::P<sub>PRM1</sub>-mCherry::hygB stl1::P<sub>STL1</sub>-YFP::URA3 bmh2::P<sub>BMH2</sub>-CFP::TRP1* | 26 |
| RB3691 | *MATa bar1Δ ssk22Δ::kanMX prm1::P<sub>PRM1</sub>-mCherry::hygB stl1::P<sub>STL1</sub>-YFP::URA3 bmh2::P<sub>BMH2</sub>-CFP::TRP1* | 26 |
| RB3695 | *MATa bar1Δ ssk1Δ::hisMX ssk22Δ::kanMX prm1::P<sub>PRM1</sub>-mCherry::hygB stl1::P<sub>STL1</sub>-YFP::URA3 bmh2::P<sub>BMH2</sub>-CFP::TRP1* | 26 |
| RB3376a | *MATa bar1Δ slt2Δ::kanMX prm1::P<sub>PRM1</sub>-mCherry::hygB stl1::P<sub>STL1</sub>-YFP::URA3 bmh2::P<sub>BMH2</sub>-CFP::TRP1* | 26 |
| RB3396a | *MATa bar1Δ fps1Δ::his prm1::P<sub>PRM1</sub>-mCherry::hygB stl1::P<sub>STL1</sub>-YFP::URA3 bmh2::P<sub>BMH2</sub>-CFP::TRP1* | 26 |
| RB3710 | *MATa bar1Δ rgc1Δ::kanMX prm1::P<sub>PRM1</sub>-mCherry::hygB stl1::P<sub>STL1</sub>-YFP::URA3 bmh2::P<sub>BMH2</sub>-CFP::TRP1* | 26 |
| RB3717 | *MATa bar1Δ rgc2Δ::kanMX prm1::P<sub>PRM1</sub>-mCherry::hygB stl1::P<sub>STL1</sub>-YFP::URA3 bmh2::P<sub>BMH2</sub>-CFP::TRP1* | 26 |
| RB3722 | *MATa bar1Δ rgc1Δ::kanMX rgc2Δ::hisMX prm1::P<sub>PRM1</sub>-mCherry::hygB stl1::P<sub>STL1</sub>-YFP::URA3 bmh2::P<sub>BMH2</sub>-CFP::TRP1* | 26 |
| RB3937 | *MATa bar1Δ hog1T100A::hog1 prm1::P<sub>PRM1</sub>-mCherry::hygB stl1::P<sub>STL1</sub>-YFP::URA3 bmh2::P<sub>BMH2</sub>-CFP::TRP1* | This study |
| RB3938 | *MATa bar1Δ hog1T100M::hog1 prm1::P<sub>PRM1</sub>-mCherry::hygB stl1::P<sub>STL1</sub>-YFP::URA3 bmh2::P<sub>BMH2</sub>-CFP::TRP1* | This study |
| YGB5938 | *MATa bar1Δ prm1::P<sub>PRM1</sub>-mCherry::hygB stl1::P<sub>STL1</sub>-YFP::URA3 stl1::P<sub>STL1</sub>-tdTomato::LEU2* | This study |
| YRB3801 | *MATa bar1Δ hog1Δ::kanMX prm1::P<sub>PRM1</sub>-mCherry::hygB stl1::P<sub>STL1</sub>-YFP::URA3 bmh2::P<sub>BMH2</sub>-CFP::TRP1* | 26 |

## Author contributions

PD performed most of the experiments. RB did the initial experiments and strain construction. AC measured glycerol. JC performed western blots. DS made strains. PD, RB, JC and DS, together with ACL analyzed the data. ACL wrote the paper.

## Acknowledgements

We thank Jeremy Thorner for plasmid pJT3557. We thank Gustavo Vasen, Pablo Aguilar and Andreas Constantinou for their comments on the manuscript. This work was supported by grants from the Argentine Agency of Research and Technology (PICT2010-2248, PICT2013-2210 and PICT2015-3824) to ACL.

## Additional Information

The authors declare no competing interests

